# Survey of activation-induced genome architecture reveals a novel enhancer of *Myc*

**DOI:** 10.1101/2022.09.14.508046

**Authors:** Wing Fuk Chan, Hannah D Coughlan, Michelle Ruhle, Nadia Iannarella, Carolina Alvarado, Joanna R. Groom, Christine R Keenan, Andrew J. Kueh, Adam K. Wheatley, Gordon K Smyth, Rhys S Allan, Timothy M Johanson

## Abstract

The transcription factor Myc is critically important in driving cell proliferation, a function that is frequently dysregulated in cancer. To avoid this dysregulation Myc is tightly controlled by numerous layers of regulation. One such layer is the use of distal regulatory enhancers to drive *Myc* expression. Here, using chromosome conformation capture to examine B cells of the immune system in the first hours after their activation, we reveal a previously unidentified enhancer of myc. The interactivity of this enhancer coincides with a dramatic, but discrete, spike in *Myc* expression 3 hours post-activation. However, genetic deletion of this region, has little impact on *Myc* expression, Myc protein level or *in vitro* and *in vivo* cell proliferation. Examination of the enhancer deleted regulatory landscape suggests that enhancer redundancy likely sustains *Myc* expression. This work highlights not only the importance of temporally examining enhancers, but also the complexity and dynamics of the regulation of critical genes such as *Myc*.

## INTRODUCTION

In order to mount an appropriate immune response, immune cells process activation signals from pathogens and immune accessory cells. The sum of these signals induce proliferation, differentiation and in the case of lymphocytes, clonal expansion. The magnitude of this response is critical. An underestimation of the threat could be fatal, while an overestimation of the danger could lead to autoimmune damage. Thus, activation induces dramatic and rapid, but tightly controlled molecular changes in immune cells, including transcriptional^1^, epigenetic^2,3^, proteomic^4^, metabolomic^5^ and three-dimensional genome organisational changes^1^.

Among the most archetypal and well-studied of these changes are those that occur to the gene of the transcription factor *Myc* and its protein product^6^. Upon lymphocyte activation Myc is rapidly upregulated in a highly controlled manner in order to oversee further transcriptional changes central to immune cell activation^7–9^.

Unsurprisingly, given its importance in both the immune context and in all healthy and diseased cell proliferation^10^, the body of research on the regulation of *Myc* is extremely rich^11,12^. Interestingly, many of the fundamental discoveries involving the regulation of *Myc* were made in immune cells. For example, shortly after the first reports of the existence and function of distal regulatory enhancers^13^, *Myc* was among the first eukaryotic genes in which regulation by these gene regulatory elements was explored^14,15^. While these early studies explored the role of aberrant enhancement of *Myc* expression driven largely by translocation, the normal genomic location of *Myc,* within a large gene-poor region in both mice and humans, made it an excellent model for the exploration of distal gene regulation. A large fraction of this regulation occurs in three-dimensions, with distal enhancers being physically drawn to the *Myc* promoter to drive expression. Thus, it is unsurprising that the invention of chromosome conformation capture, which can reveal the three-dimensional proximity of DNA to other DNA^16,17^, drove significant advancement of our understanding of the distal regulation of *Myc*^11^. These studies revealed an elaborate, cell-type- and cell-state-specific three-dimensional enhancer landscape controlled by umpteen transcription factors, histone modifications, DNA methylation, long non-coding RNAs^18^, amongst other mechanisms^*11*^.

Here, via exploring changes in three-dimensional genome organisation in the first hours post-B cell activation, we reveal a previously uncharacterised upstream enhancer of *Myc,* apparent for mere hours following activation, which accompanies the rapid and dramatic spike in post-activation *Myc* expression. However, genetic removal of this enhancer leads to minimal impact on either *Myc* expression or Myc protein level, likely due to differential enhancer usage or enhancer redundancy sustaining critical levels of Myc.

## RESULTS

### Activation induces genome reorganisation upstream of *Myc* promoter

To begin exploring the three-dimensional genome architectural change that may regulate early activation-induced transcriptional change, we examined paired chromosome conformation capture (*in situ* HiC) and RNA-Seq data of activation-induced B cell differentiation^1^ with a particular focus on the changes between naïve and 3 hour activated B cells (Figure 1A).

**Figure 1:**
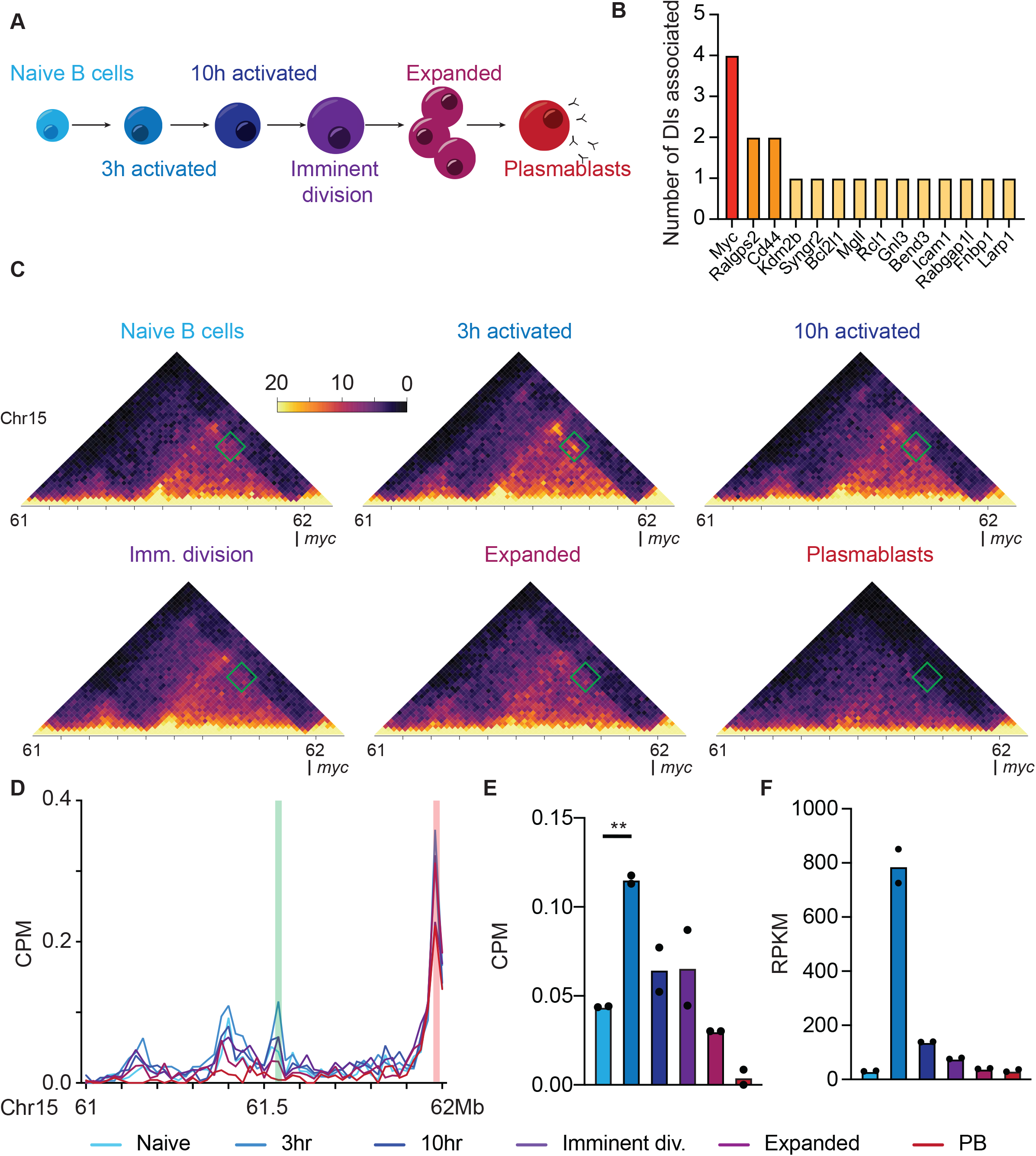
Activation-induced genome reorganisation upstream of *Myc* promoter. **A)** Schematic of the stages of B cell differentiation examined using *in situ* HiC and RNA-Sequencing. **B)** Number of differential interactions (DIs) associated with all DI-associated differentially expressed genes (DEs) in the top 100 DEs between naïve and 3 hour activated B cells. **C)** Normalised *in situ* HiC contact matrices at 20kbp resolution showing genomic region upstream of the *Myc* gene promoter of all six stages of B cell differentiation. Region shown chr15:61-62.1Mb. Colour scale indicates the normalised counts per bin pair. Green box highlights 3 hour-specific interacting region of interest. **D)** Virtual 4C of the same region of chromosome 15 shown in C) at 20kbp resolution from viewpoint of the myc promoter (chr15:61983391-61990390) plotted as counts per million (CPM) in all six stages of B cell differentiation. Red highlight is the viewpoint. Green highlight in the 3 hour specific interactive region of interest. Mean of duplicate biological replicate samples shown. **E)** CPM of interaction between the *Myc* promoter and the 3 hour specific interactive region of interest across all stages of B cell differentiation. Unpaired two-tailed T test used to test for significance. ** denotes P<0.005. **F)** Reads per million per kilobase of *Myc* transcripts across all stages of B cell differentiation derived from RNA-Sequencing.

First, we examined all the differential interactions (DIs) between naïve and 3 hour activated B cells associated with the top 100 differentially expressed genes (DEs) between the same cells (Supplementary Table 1). DIs are statistically significant differences in the DNA-DNA interaction frequency between any two points on the linear genome between samples, determined using diffHiC^19^, and denote changes in enhancer-promoter DNA loops, or other genome architectural changes. The top 100 DEs associate with just 19 DIs, four of which associate with the promoter of the *Myc* gene (Figure 1B). This suggests that *Myc* is one of the few genes regulated by early activation-induced genome architectural change in B cells.

To further define the nature and position of the *Myc* associated activation-induced genome architectural changes, we generated contact matrices of the genomic regions containing the DIs (Figure 1C) and performed virtual 4C of the same region using the *Myc* promoter (2kbp upstream and 5 kbp downstream) as the viewpoint (Figure 1D, viewpoint shown in red). This analysis revealed a previously unknown DNA-DNA interaction between the *Myc* promoter and an upstream region (Figure 1C,D in green). This statistically significant change in interactivity (*P*=0.001, unpaired twotailed t-test,) appears to be highly 3-hour specific (Figure 1E) and is the strongest architectural change in the region (Supplementary Figure 1) between naïve and 3 hour activated B cells. Interestingly, the interactivity of this region and the *Myc* promoter (Figure 1E) across activation-induced B cell differentiation is reflective of the expression pattern of *Myc* in the same cells (Figure 1F). This may reflect a regulatory role for the region in the expression of *Myc.*

### Deletion of one putative enhancer of *Myc* alters *Myc* expression

To explore potential gene regulatory roles for the putative activation-induced enhancers discovered upstream of *Myc*, we first sought to clarify the potential functions of the regions using available epigenetic data. Overlaying our *in situ* HiC data with publicly available ATAC-Sequencing^20^ and H3K27 acetylation and H3K4 mono-methylation chromatin immunoprecipitation data^21^ (generally associated with DNA accessibility, active enhancers and active/primed enhancers, respectively), we revealed part of the epigenetic landscape of the region in naïve B cells (Figure 2A). This analysis highlighted four genomic regions of particular interest (Region 1-4). Region 1,2 and 3 (R1-3) were both accessible and contained epigenetic marks consistent with enhancers. Region 4 was accessible and in contact with the *Myc* promoter, but contained no detectable H3K27ac or H3K4me1 modifications.

**Figure 2:**
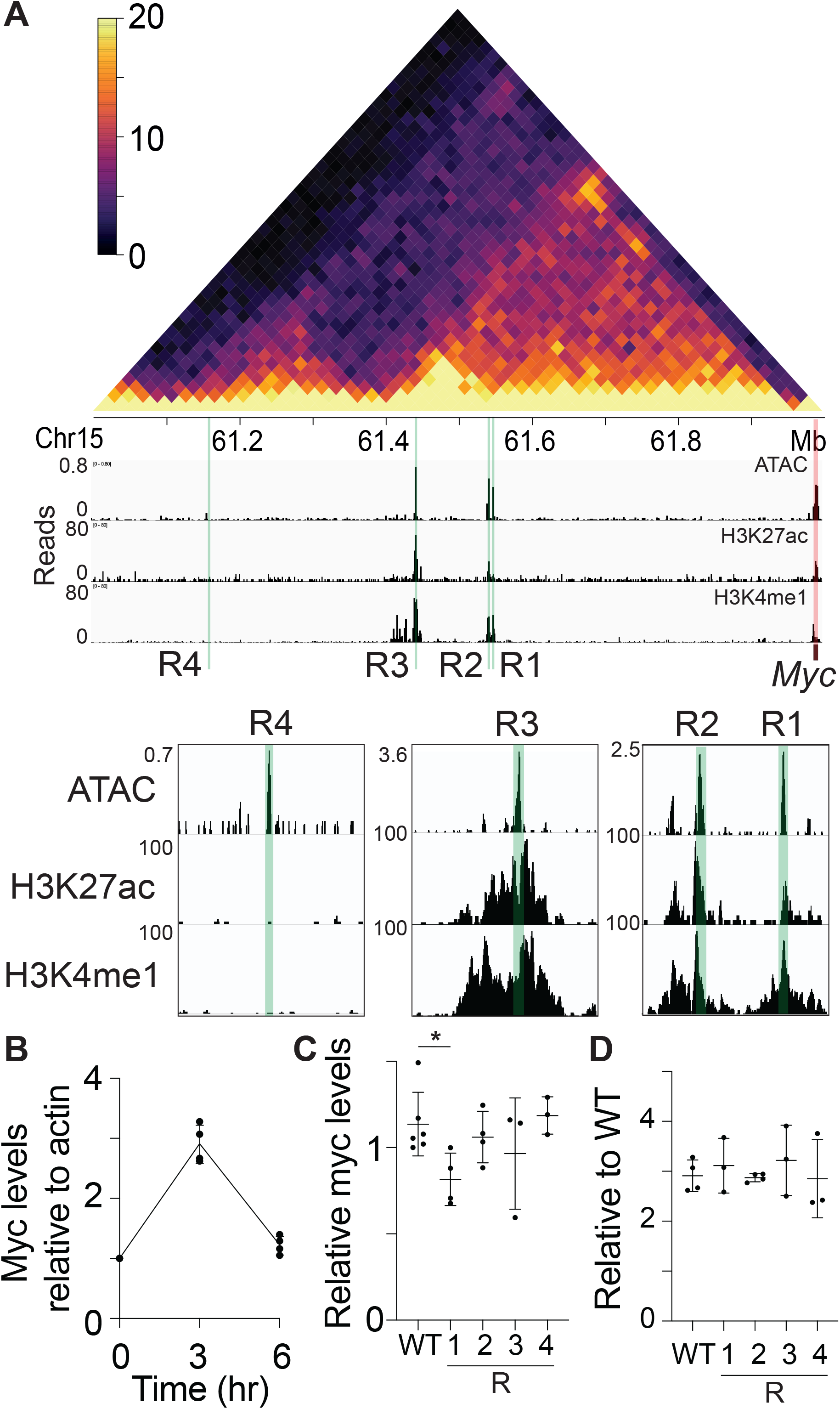
Genetic deletion of putative enhancer regions upstream of *Myc* reveals a novel enhancer. Overlay of *in situ* HiC (top panel) from 3 hour activated B cells at 20 kbp resolution and DNA accessibility (ATAC-sequencing)^20^, H3K27 acetylation (H3K27ac)^21^ and H3K4 mono-methylation (H3K4me1)^21^ from naïve B cells upstream of the *Myc* promoter (chr15:61-62Mb). Color scale indicates the normalised counts per bin pair. Regions 1-4 (R1-4) are highlighted in green. The *Myc* promoter is highlighted in red. Insets show the same data at regions of interest at higher resolution. **B)** Quantitative RT-PCR of *Myc* transcript levels relative to *Actb* transcript (encoding β-actin) in A20 cell line 3 or 6 hours post-activation with lipopolysaccharide (LPS), relative to 0 hours. Data representative of 3 independent experiments and normalised to β-actin. Mean +/− SD shown. **C)** Quantitative RT-PCR of *Myc* transcript levels relative to *Actb* transcript in A20 cells with either region 1, 2, 3 or 4 (R1-4) genetically deleted. Region 1-4 data presented as relative to the wild-type (WT) A20 sample. * denotes P<0.05. Data representative of at least 3 independent experiments and normalised to *Actb.* Mean +/− SD shown. **D)** Quantitative RT-PCR of *Myc* transcript levels in A20 cells activated for 3 hours with LPS with either region 1, 2, 3 or 4 (R1-4) genetically deleted. Region 1-4 data presented as relative to the levels prior to activation. Data representative of at least three independent experiments and normalised to *Actb.* Mean +/- SD shown.

To definitively characterise the function of these regions in regulating *Myc* expression, we genetically removed each region using clustered regularly interspaced short palindromic repeats (CRISPR)/Cas9 technologies in the A20 B cell line (Supplementary Fig 2A). First, we confirmed that activation induced *Myc* expression shows a similar pattern in the A20 cell line as in primary B cells (Figure 2B). We then quantified the impact of the genetic removal of regions 1, 2, 3 or 4 (Supplementary Figure 2A) on *Myc* expression levels in the steady state (Figure 2C) or 3 hours after activation (Figure 2D) using qRT-PCR. Only deletion of region 1 in the steady state has any impact on the levels of *Myc,* inducing a small but significant reduction (Figure 2C). Interestingly, this reduction is no longer present in the same cells activated for 3 hours (Figure 2D), suggesting different mechanisms function to regulate *Myc* expression after activation. Regardless, the impact on *Myc* expression upon genetic deletion of region 1 demonstrated that this region functions as an enhancer of *Myc.* To more fully explore its function, we generated a mouse strain with this region deleted (Supplementary Figure 2B,C), termed Myce (*Myc* enhancer deleted) mice.

### Myce mice have normal *in vitro* B cell activation responses

After confirming that Myce mice have a normal immune compartment in various organs (Supplementary Figure 3 A-C), we performed an *in vitro* B cell activation assay to quantify activation induced proliferation and survival. In brief, B cells were isolated from Myce mice or wild-type littermate controls, activated with lipopolysaccharide (LPS) then their Myc protein levels, proliferation and death were observed over time. These experiments revealed little to no differences between Myce mice and littermate control B cells in either the levels of Myc protein in the early hours after activation (Figure 3A, Supplementary Figure 3D), total cell number in culture over their 3-day activation-induced expansion (Figure 3B), or the B cells survival (Figure 3C) or division rate (Figure 3D) during this culture period. Similar results were seen in T cell activation cultures (Supplementary Figure 3E-H). These experiments emphatically show that the deletion of the enhancer of *Myc* has no significant impact on either the activation-induced levels of Myc, nor the B cell response to *in vitro* activation.

**Figure 3:**
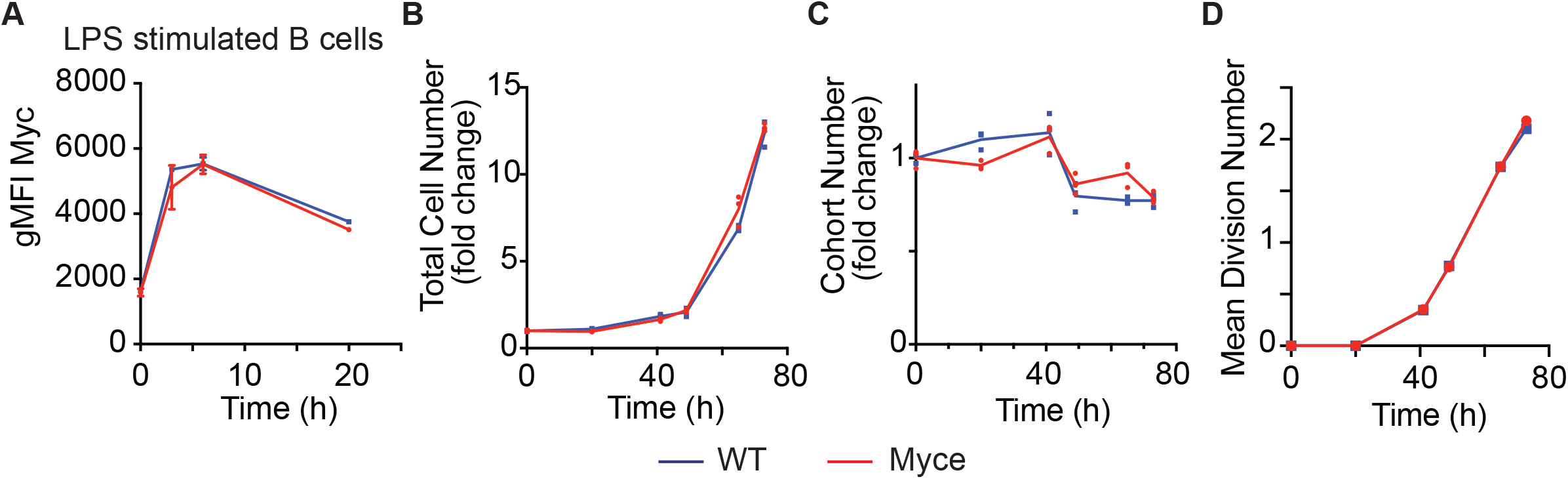
Myce mice have normal *in vitro* B cell activation responses. **A)** Plot of geometric mean fluorescence intensity (gMFI) of Myc protein detected by flow cytometry in Myce mice or wild-type littermate control B cells at 0, 3, 6 or 20 hours postactivation with LPS. **B)** Fold change in total B cell number relative to 0 hours across 72 hours post-activation with LPS in Myce mice and wild-type littermate controls. **C)** Cohort number (number of B cells in each division divided by 2^division number) as a measure of B cell survival over 72 hours post-activation with LPS in Myce mice and wild-type littermate control B cells. **D)** Mean division number, measured by Cell Trace Violet division tracker, over 72 hours post-activation with LPS in Myce mice and wild-type littermate control B cells. All data is representative of two independent experiments. Mean +/− SEM shown.

### Myce mice have normal *in vivo* B cell activation responses

We next determined the impact of the deletion of the *Myc* enhancer on *in vivo* immune responses. First, we infected Myce mice and wild-type littermate controls with influenza (X31). At the peak of the infection (day 8) we examined total cell number and B cell number in the bronchoalveolar lavage (Figure 4A) and mediastinal lymph nodes (Figure 4B) of infected Myce mice and littermate controls (as well as uninfected littermate controls). This enumeration revealed no significant differences in lung infiltrate between the Myce mice and controls. Similarly, other immune cell types, both antigen specific and not, were not significantly affected by the enhancer deletion (Supplementary Figure 4A-B).

**Figure 4:**
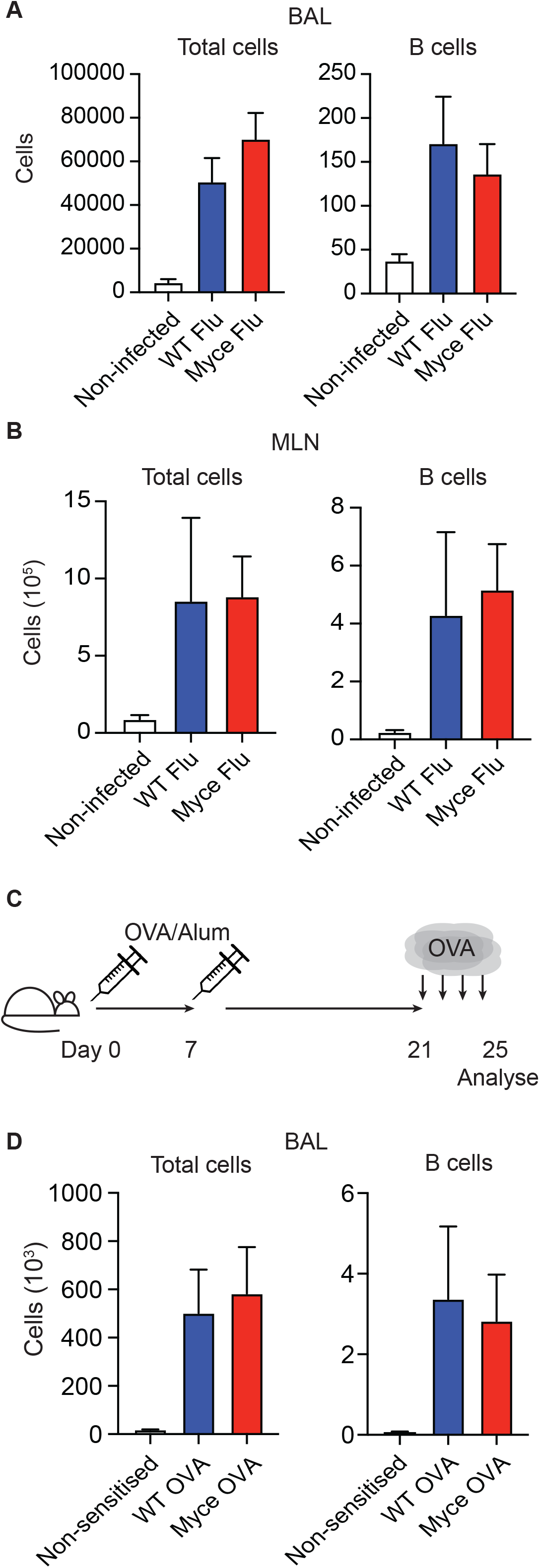
Myce mice have normal *in vivo* B cell activation responses. **A)** Number of total live cells and B cells detected in the bronchoalveolar lavage (BAL) of Myce mice, littermate controls (WT Flu) or mice not infected with influenza (Non-infected) at Day 8 post-infection. **B)** Number of total live cells and B cells detected in the mediastinal lymph nodes (MLN) of Myce mice, littermate controls (WT Flu) or mice not infected with influenza (Non-infected) at Day 8 post-infection. **C)** Schematic of the experimental setup for the model of asthma using OVA/alum sensitisation. In brief, mice are sensitised with OVA and aluminium hydroxide on Day 0 and 7, rested for 14 days then challenged with nebulized OVA for 4 consecutive days. Analysis is performed one day after the final OVA challenge. **D)** Number of total live cells and B cells detected in the bronchoalveolar lavage (BAL) of Myce mice or littermate controls (WT OVA) sensitised with OVA/Alum or littermate controls sensitised with Alum alone (Non-sensitised). Data derived from at least two independent experiments. Mean+/− SEM shown.

We also tested the immune response generated in allergic lung inflammation using intraperitoneal ovalbumin sensitization followed by nebulised OVA challenge (Figure 4C)^22^. Cell populations in the bronchoalveolar lavage (BAL) were examined the day after the final OVA challenge. Enumeration of total cells, B cells (Figure 4D) and other immune cell types (Supplementary Figure 4C) revealed no significant differences between Myce mice and littermate controls.

Together with the unchanged numbers or proportions of immune cells in steady state Myce mice (Supplementary Fig3A-C), these experiments suggest that the deletion of myce has little to no impact on the maintenance or activation response of immune cells *in vivo.*

### Myce mice cells display altered chromatin interactions with the *Myc* promoter

We next sought to determine why deletion of the *Myc* enhancer had little to no impact on either *Myc* expression, Myc protein levels or immune responses in cells from the Myce mice. To do so, we examined the consequence of the deletion on the genome architecture upstream of *Myc* in both the Myce mice and littermate controls. In brief, we isolated B cells from either Myce mice or wild-type littermate controls, activated them *in vitro* for 3 hours then sorted the activated and naïve B cells from the culture. A proximity ligation protocol followed by DNA precipitation was then performed on these cells, before qPCR was used to determine the frequency of DNA-DNA interaction between the *Myc* promoter and a number of upstream genomic regions.

The frequency of interaction determined by this qPCR-based method correlated well with *in situ* HiC data from the same region (Figure 5A–C) with regions of high or low interactivity detected similarly by both methods. Importantly, it also revealed that in 3 hour activated B cells the interaction frequency at each examined region upstream of the deleted enhancer is greater in the Myce mice B cells than littermate controls (Figure 5B left of the green dotted line). This is in contrast to the regions downstream of the deletion in both Myce mice and littermate controls (Figure 5B, right of the green dotted line) and in naïve B cells either up or downstream (Figure 5C), suggesting the change in interactivity is deletion and activation induced. This change in interactivity can be more clearly observed in the frequency of interaction in the activated Myce B cells relative to the WT (Figure 5D). Furthermore, summing all relative threshold cycle (Ct) values upstream or downstream of the enhancer deleted region in both naïve and activated B cells reveals a significant (*P* = 0.04, paired twotailed t-test) increase in interaction frequency upstream of the deletion, specifically in activated B cells (Figure 5E).

**Figure 5:**
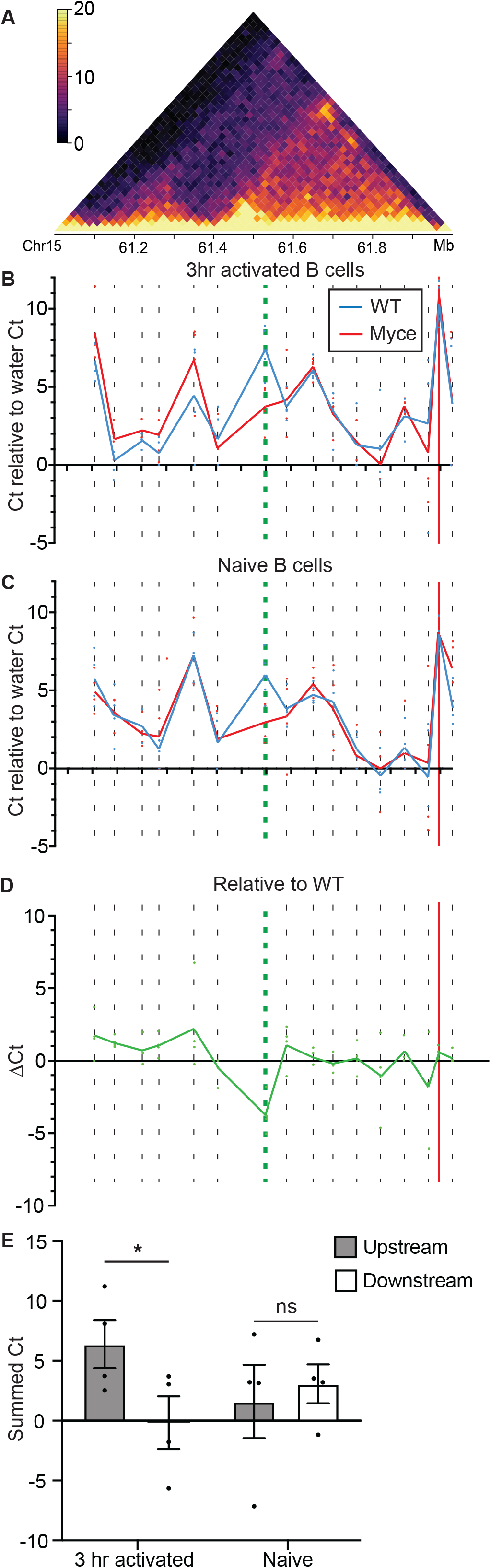
Altered interactions with the *Myc* promoter in Myce mice B cells. **A)** *In situ* HiC contact matrix at 20 kbp resolution showing upstream of the *Myc* promoter (chr15:61-62Mb) in 3 hour activated B cells. Color scale indicates the normalised counts per bin pair. **B,C)** Threshold cycle (Ct) from qPCR targeting the *Myc* promoter and its proximity-ligated DNA partners from across the region upstream of the promoter (Supplementary Table 2), relative to a water control Ct, from **B)** 3 hour activated (CD69^+^) or **C)** naïve (CD69^-^) Myce mice or wild-type littermate control B cells. Dotted lines represent regions targeted for amplification by qPCR. Green dotted line represents primer targeting enhancer deleted region. Red vertical line represents *Myc* promoter. Mean of four independent experiments shown. **D)** Threshold cycle (Ct) from qPCR targeting the *Myc* promoter and its proximity ligated DNA partners from across the region upstream of the promoter in activated Myce mice B cells relative to the activated wild-type littermate control Ct. Dotted lines represent regions targeted for amplification by qPCR. Green dotted line represents primer targeting enhancer deleted region. Red vertical line represents *Myc* promoter. Mean of four independent experiments shown. **E)** The sum of the Myce mice relative to littermate controls threshold cycle (myce Ct - WT Ct) from upstream and downstream of the deleted region (left and right of the green dotted line, respectively, in D) in 3 hour activated and naïve B cells. Mean +/− SEM from four independent experiments shown. * denotes P<0.05.

These experiments suggest that the reason that Myc levels, or indeed immune responses, are unchanged in the Myce mice B cells under various activation conditions is that in the absence of the enhancer the *Myc* promoter simply continues searching the upstream DNA to find another enhancer to sustain its expression. This is reflected in the increased interaction frequencies upstream of the enhancer deletion in Myce mice activated B cells.

## DISCUSSION

Here, through a comprehensive examination of genome organisational changes in the hours after B cell activation, we identified a previously undetected enhancer of *Myc,* in a highly scrutinised regulatory landscape^11^. However, genetic deletion of this enhancer has little to no impact on *Myc* expression, protein levels or *in vitro* and *in vivo* cell proliferation or survival. Further exploration revealed that *Myc* expression is maintained in the absence of this activation-induced enhancer likely via differential enhancer usage or enhancer redundancy.

Enhancer redundancy is the process by which gene expression is sustained upon the deletion of an enhancer of a gene by the action of other enhancers of the gene^23^. This process has been widely documented across the kingdoms of complex life^24–27^ and appears to be enriched in genes important in development and health, supporting robust transcription buffered against environmental and genetic perturbation. For example, the total number of predicted enhancers and the redundancy of transcription factor binding within these enhancers is predictive of a genes potential pathogenicity^28,29^. Experimentally testing this potential is challenging as deletion of enhancers in these redundant regulatory landscapes frequently results in little to no phenotypic impact^23^. However, this is not true in all cases. For example, in the *Myc* regulatory landscape, deletion of eight different enhancers within 2 megabases downstream of the myc promoter each lead to significant reductions in *Myc* expression^30,31^.

Our detailed exploration of the three-dimensional organisation in the enhancer deleted genome provides interesting insights into potential mechanisms of enhancer redundancy. As such, in 3 hour activated B cells, deletion of the activation induced enhancer appears to increase DNA-DNA interactions between the *Myc* promoter and numerous upstream enhancers. Furthermore, the magnitude of these interaction changes suggests multiple new interactions per B cell. This suggests that the redundancy observed is not simply a case of the *Myc* promoter contacting and utilising the next upstream enhancer to maintain appropriate regulation, but potentially the whole upstream regulatory landscape.

Why some enhancers exhibit redundancy and others do not is unknown. What is clear is that the combined function of enhancers is extremely context specific. For example, the enhancers of *hunchback* in *D. melanogaster* will behave either additively (the induction of expression in the presence of both enhancers is the sum of both individually) or subadditively (the induction of expression in the presence of both enhancers is less than the sum of both individually) depending on the concentration of the transcription factor, Bicoid^32^. Similarly, two enhancers of *pomc* in mice have been demonstrated to function additively in adult mice neurons, but superadditively (the induction of expression in the presence of both enhancers is greater than the sum of both individually) in young mice neurons^33^. Thus, while it is clear that transcription factors can influence enhancer usage, epigenetics, the relative position of the enhancer to the promoter or other enhancers, among many other variables, likely impact enhancer redundancy. Transcription factors likely play a role in the enhancer redundancy observed in this study as the deleted enhancer contained a predicted E-Box and an NFκB binding site. NFκB has previously been shown to regulate *Myc* expression via binding near the promoter^34^, so under normal conditions our enhancer likely forms part of this NFκB -*Myc* regulatory system.

Here we have revealed a slight, but significant change in steady state *Myc* expression levels upon deletion of a novel enhancer, revealed by a detailed examination of genome organisation just hours after cell activation. However, enhancer redundancy has made it challenging to accurately dissect the normal function of this enhancer. Nonetheless, the correlation between the temporal specificity of the interaction between our novel enhancer and the *Myc* promoter and the dramatic and discrete spike in *Myc* expression at 3 hours post-activation is compelling. Thus, we propose that under normal conditions this activation-induced enhancer assists in regulating this critical and dramatic increase in *Myc* expression. The development and application of new super-resolution live imaging technologies that allow concurrent visualisation of multiple regions of DNA and associated factors at single molecule resolution^35,36^, will likely soon enable the visualisation and dissection of not only this function, but the discussed genome architecture of the enhancer deleted *Myc* regulatory landscape and definition of the factors regulating redundancy.

## FIGURE LEGENDS

**Supplementary Figure 1:**
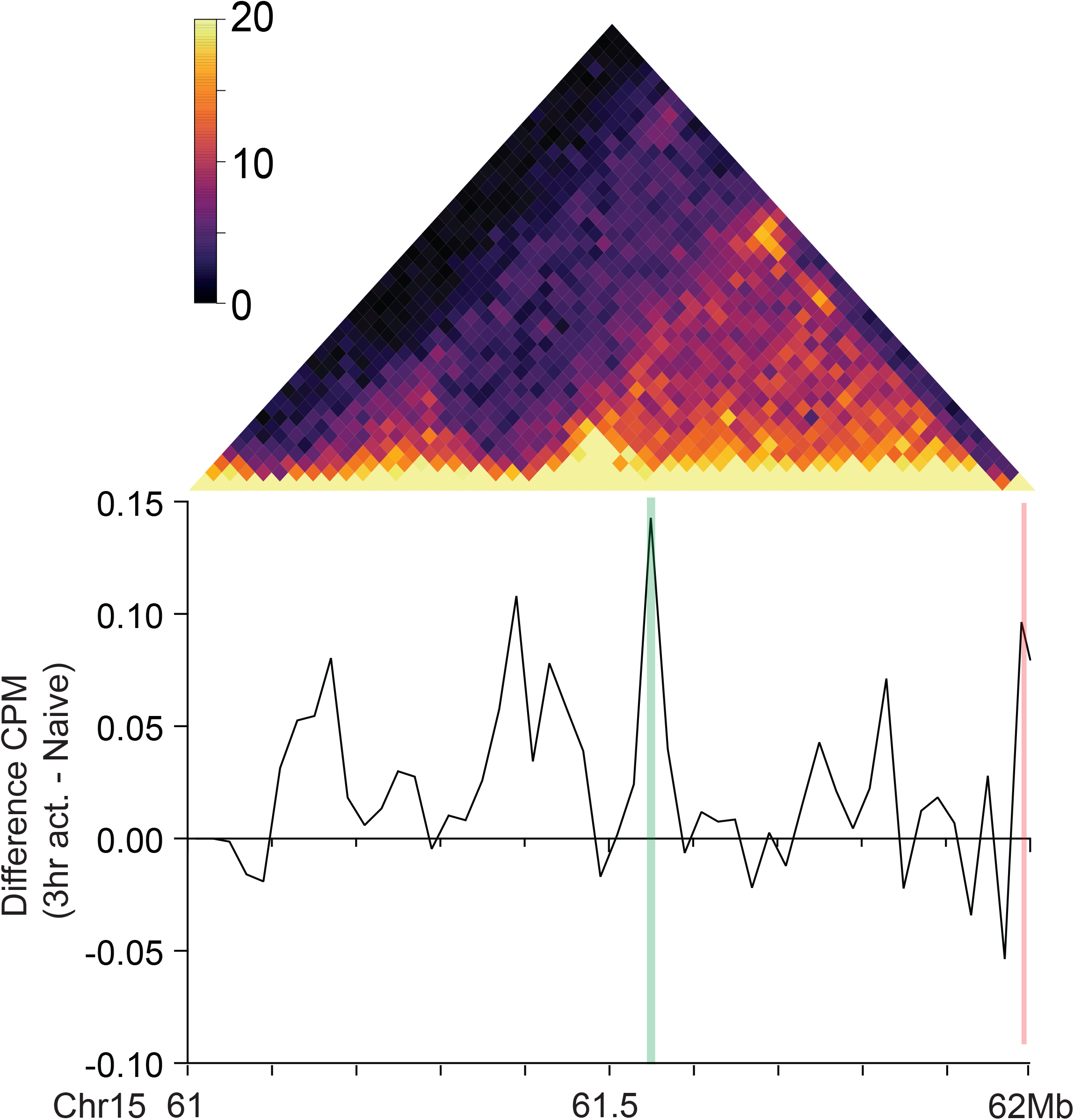
Difference between the pooled means of 3 hour activated B cell and naïve B cell *in situ* HiC libraries virtual 4C profiles (as plotted in Figure 1D) of region chr15:61-62Mb. Values are counts per million (CPM) in 20kbp bins relative to the viewpoint at the *Myc* promoter (chr15:61983391-61990390 bp) Vertical red line represents the viewpoint. Vertical green line represents the activation-induced enhancer.

**Supplementary Figure 2:**
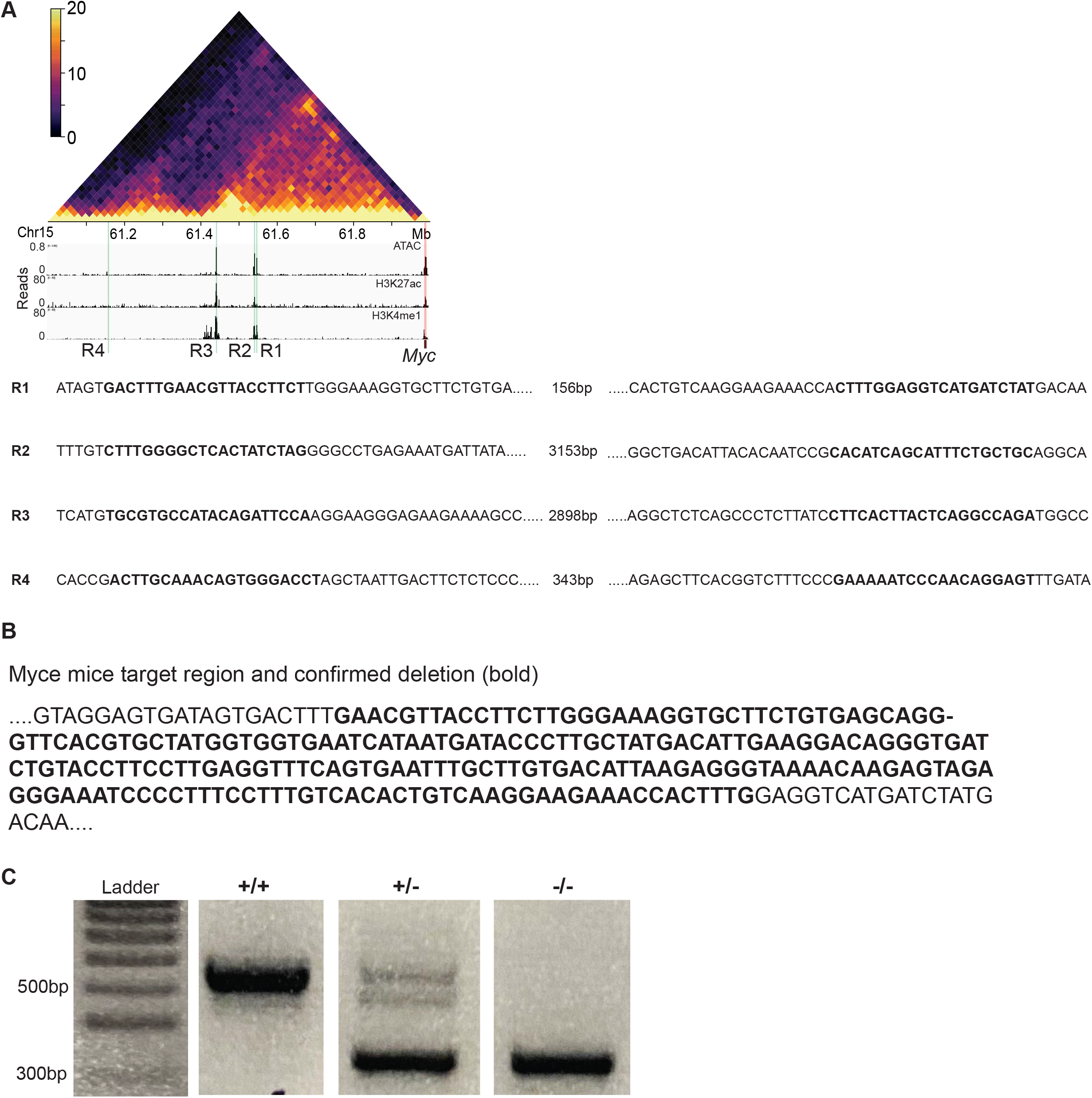
**A)** The four genomic regions (R1-4) targeted for deletion using CRISPR/Cas9 in the A20 cell line, and sequences of the Cas9 target used to induce deletion (Supplementary Table 2) **B)** The genomic region (R1) deleted in the Myce mice line. Deletion was confirmed using Sanger and Next Generation sequencing. **C)** Example of DNA band pattern derived from wild type, heterozygous and homozygous deletion of the enhancer of interest in Myce mice using PCR and gel electrophoresis. PCR primers are listed in Supplementary Table 2.

**Supplementary Figure 3:**
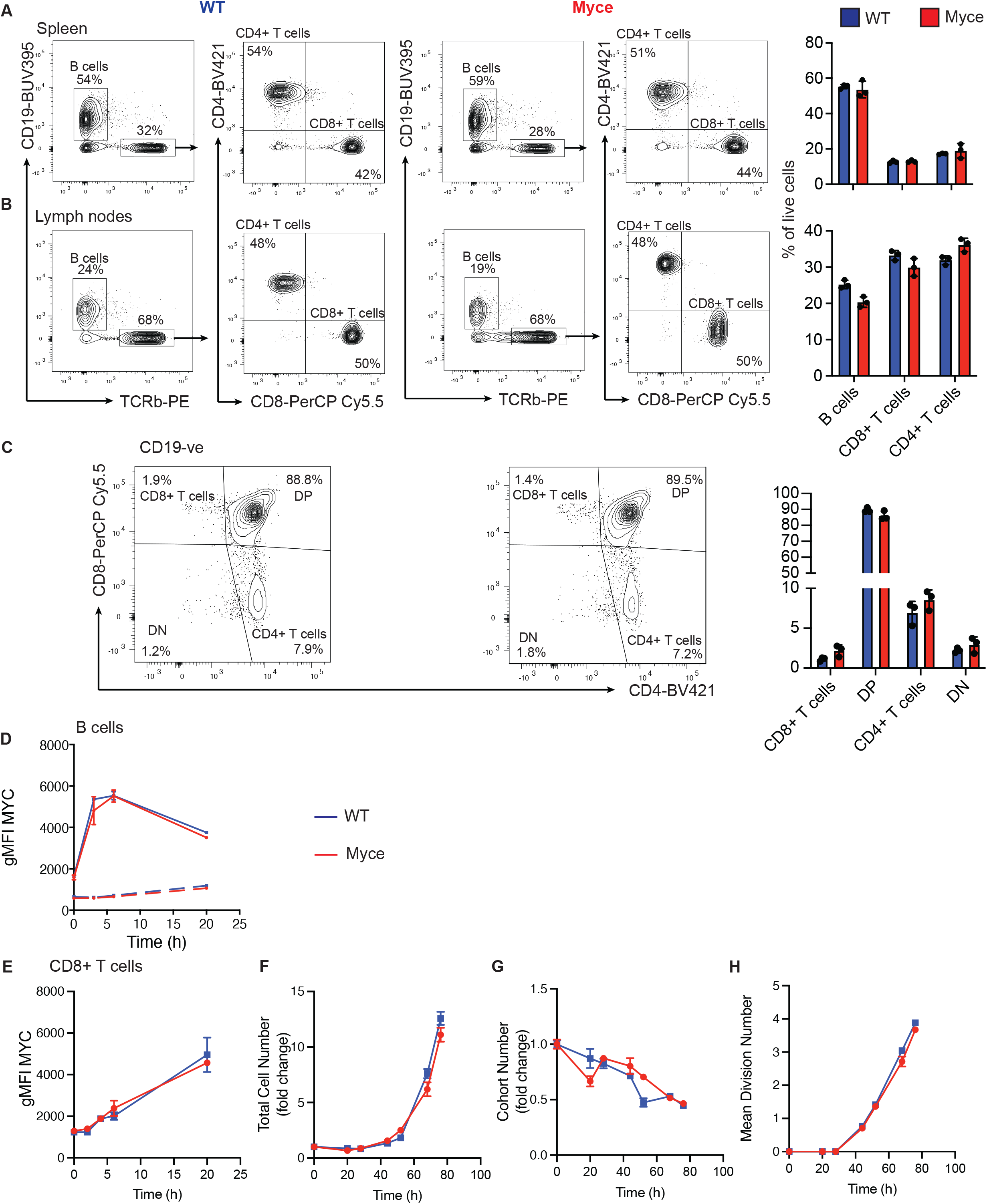
Flow cytometry profiles and quantitation of B cells and T cells in the **A)** spleen, **B)** lymph nodes and **C)** thymus of Myce mice and littermate controls. Mean+/− SD shown. **D)** The geometric mean fluorescence intensity (gMFI) of Myc staining in *in vitro* activated B cells derived from flow cytometry on Myce mice and wild-type littermate control B cells. Dotted lines indicate isotype control detection in the two populations. **E)** Plot of geometric mean fluorescence intensity (gMFI) of Myc protein in Myce mice or wild-type littermate control CD8^+^ T cells at 0, 3, 6 or 20 hours post-activation with LPS. **F)** Fold change in total CD8^+^ T cell number relative to 0 hours across 72 hours post-activation with LPS in Myce mice and wild-type littermate controls. **G)** Cohort number (number of CD8^+^ T cells in each division divided by 2^division number) as a measure of T cell survival over 72 hours post-activation with LPS in Myce mice and wild-type littermate control B cells. **H)** Mean division number, measured by Cell Trace Violet division tracker, over 72 hours postactivation with LPS in Myce mice and wild-type littermate control CD8^+^ T cells. All data is representative of two independent experiments. Mean +/− SEM shown.

**Supplementary Figure 4:**
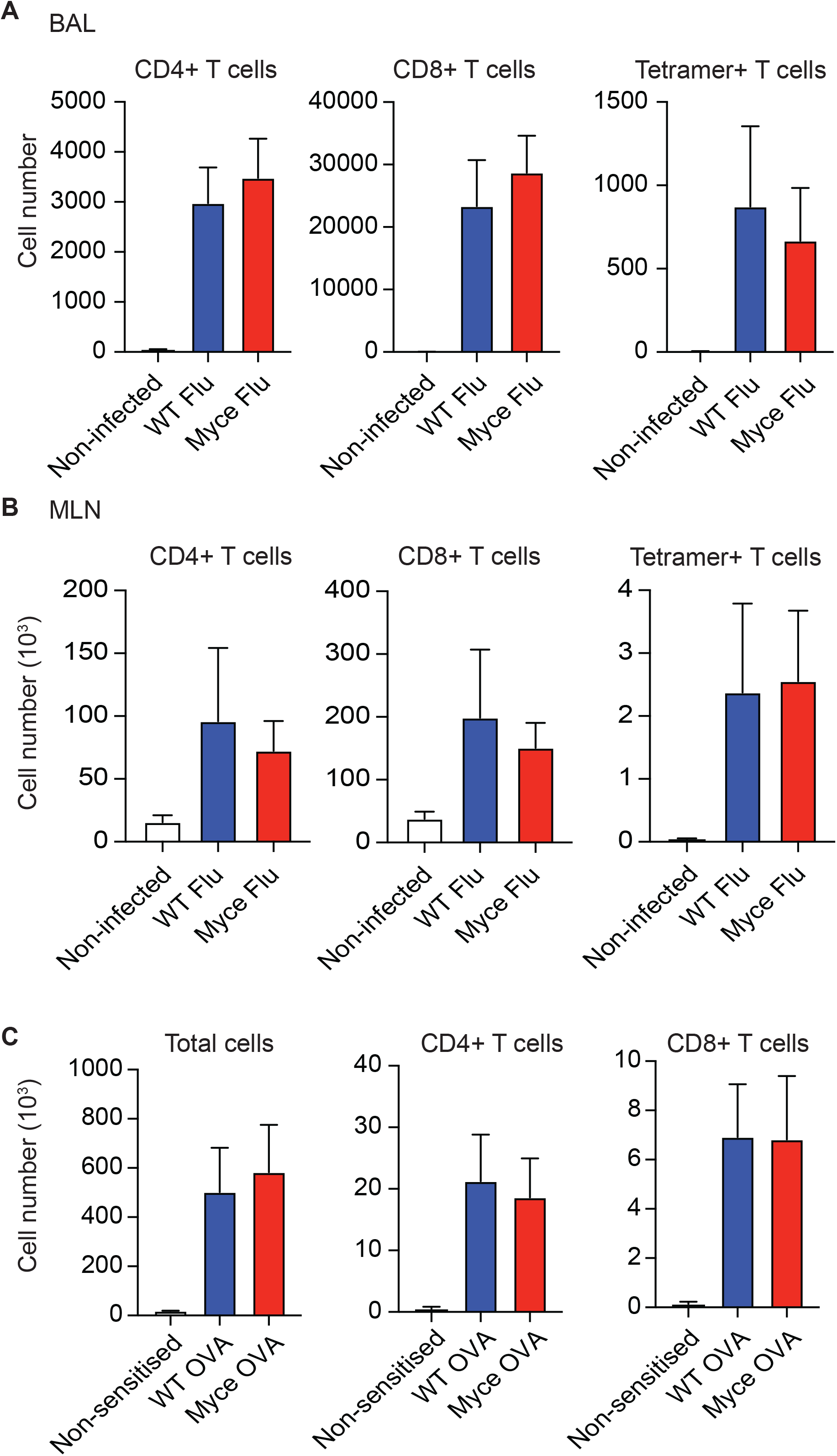
Number of CD4+, CD8+ and tetramer positive T cells detected in the **A)** bronchoalveolar lavage (BAL) or **B)** mediastinal lymph nodes (MLN) of Myce mice and littermate controls infected with influenza (Influenza A/H3N2/X31at 1×10^4^pfu) and non-infected controls at Day 8 post-infection. **C)** Number of total live cells and T cells detected in the bronchoalveolar lavage (BAL) of Myce mice or littermate controls (WT OVA) sensitised with OVA/Alum or littermate controls sensitised with Alum alone (Non-sensitised). Data derived from at least two independent experiments. Mean+/− EM shown.

**Supplementary Table 1.**
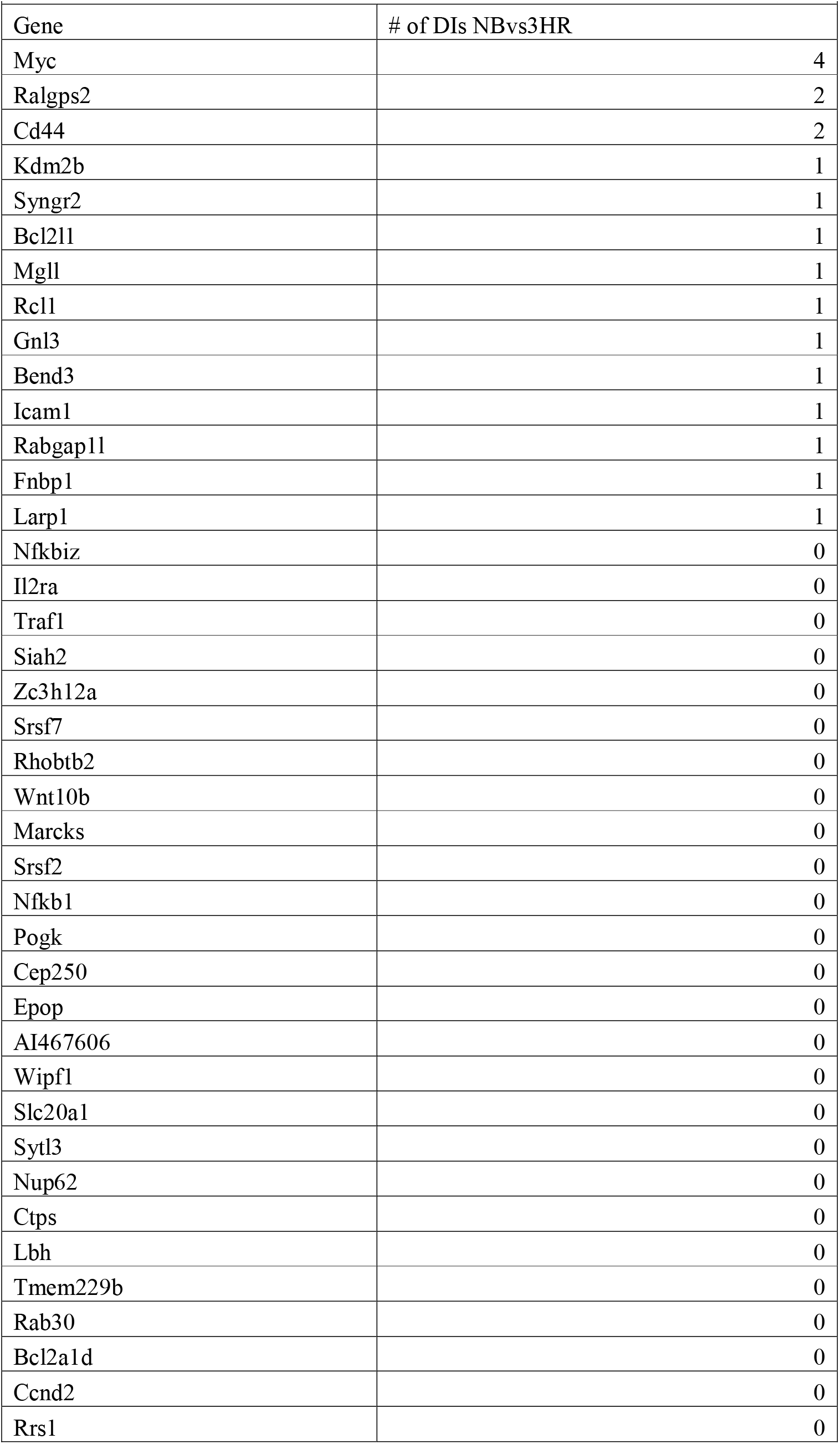
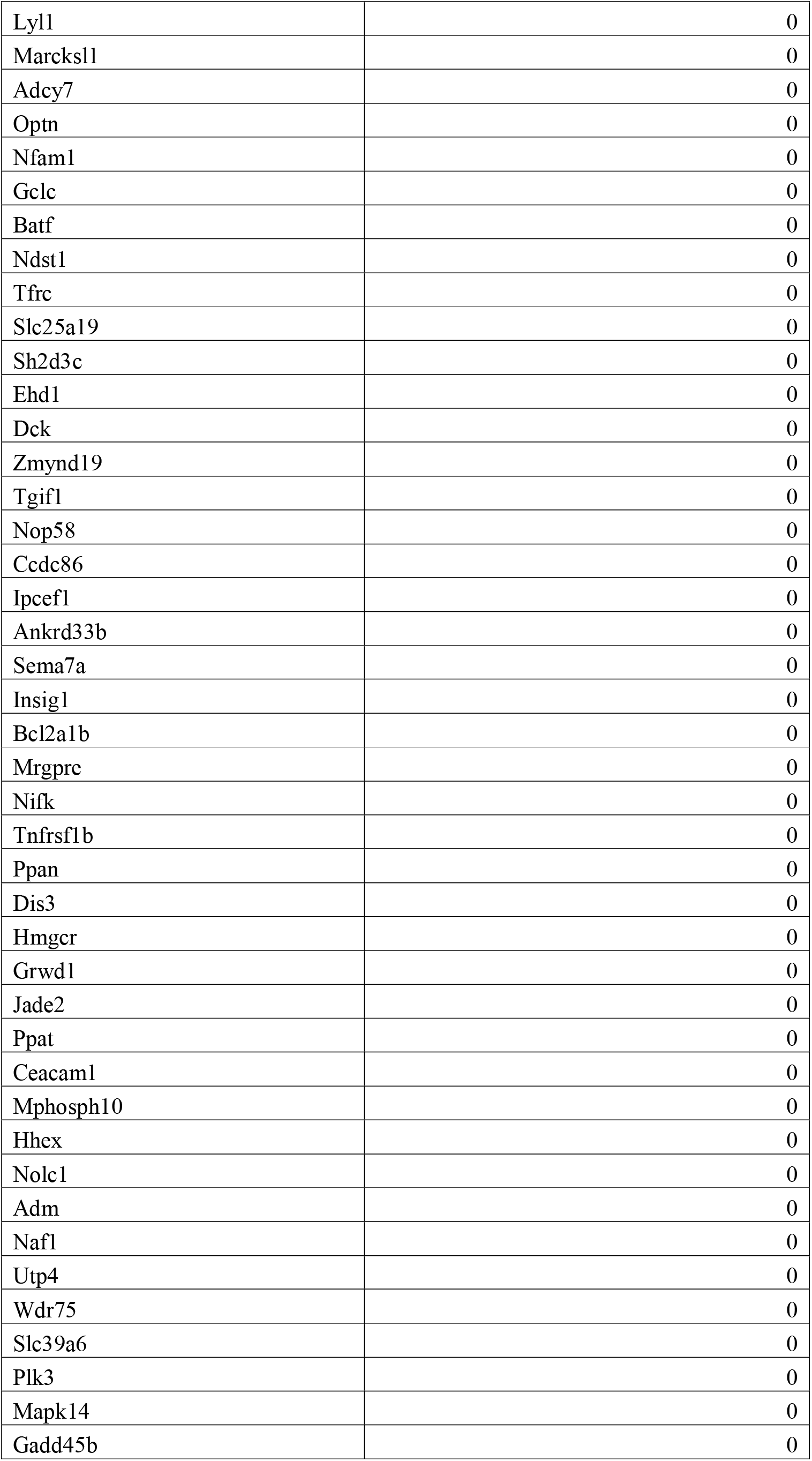
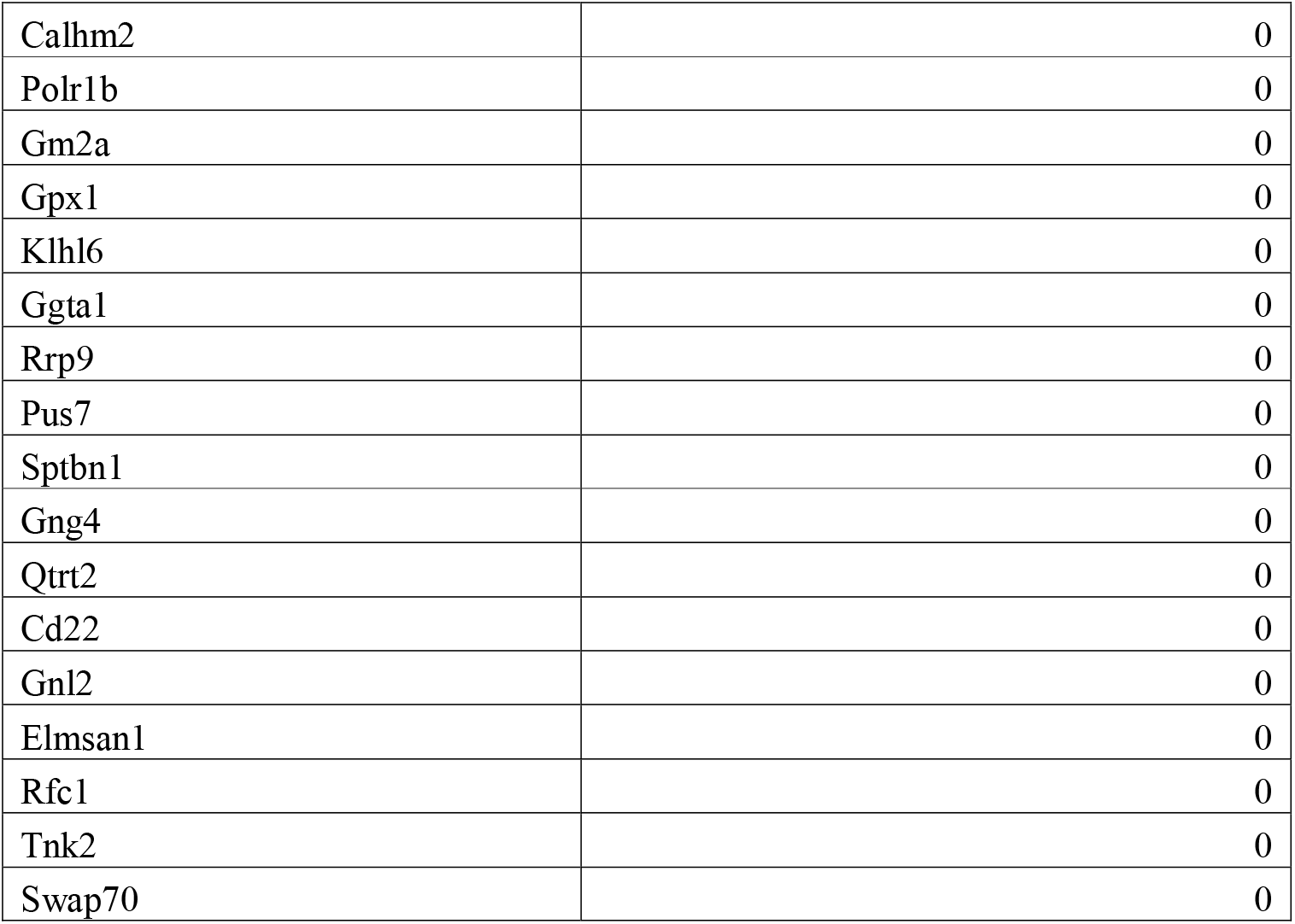
Top 100 differentially expressed genes between naïve and 3 hour activated B cells and their associated differential interactions

**Supplementary Table 2.**
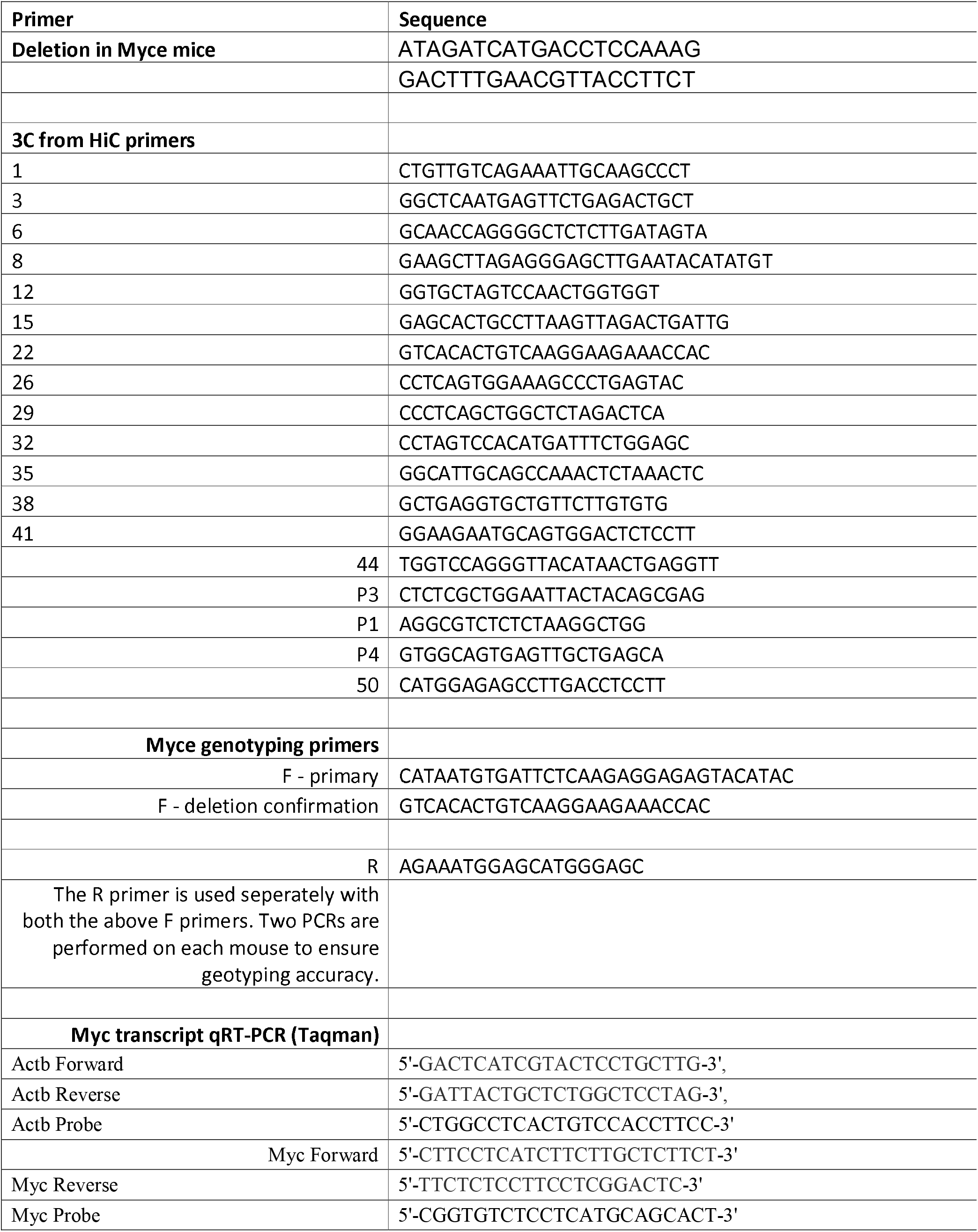
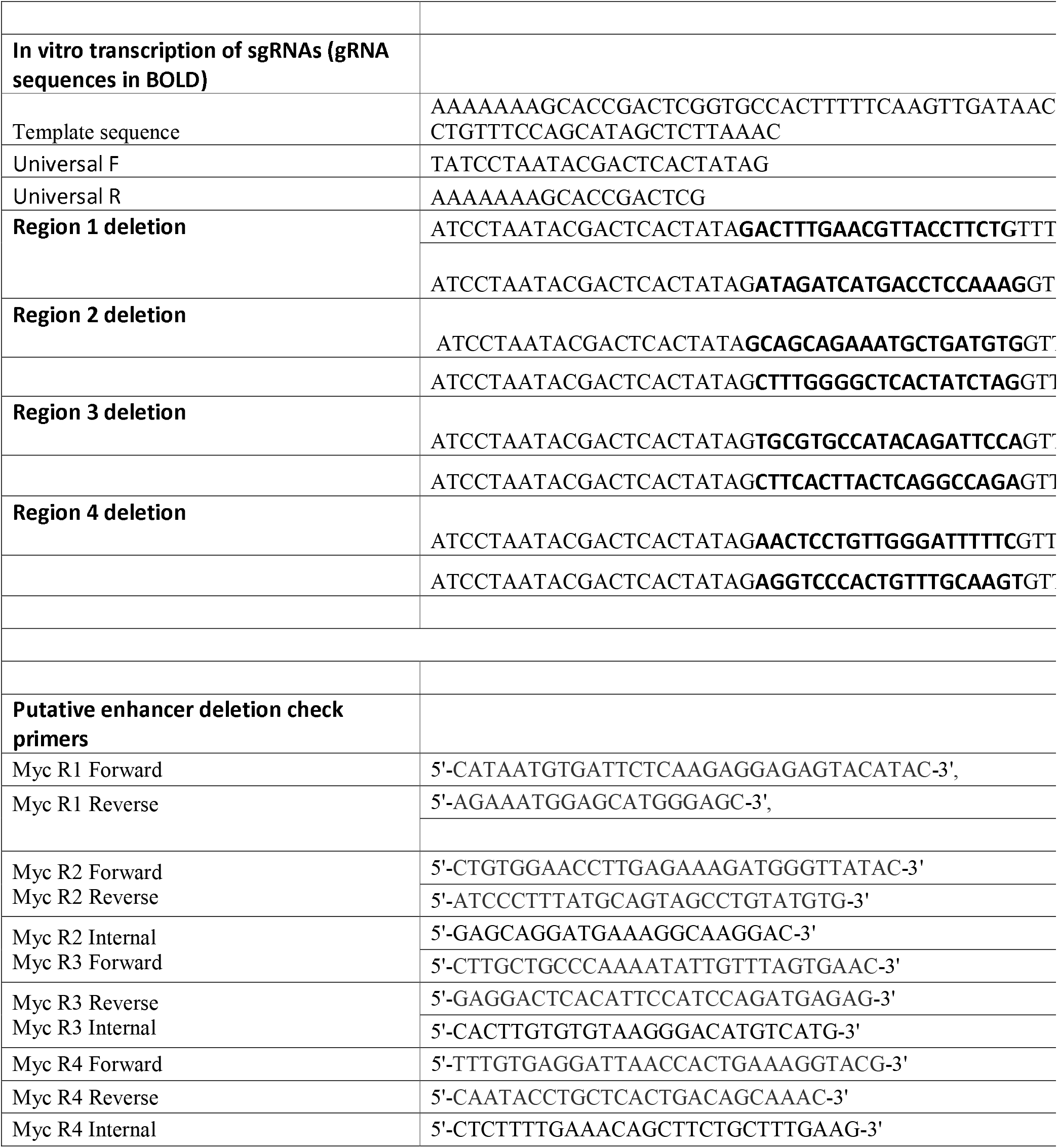
Primers used in the study

**Supplementary Table 3.** Antibodies used in this study

| **Supplementary Table 3**<br>**Antibodies used in this study** |  |  |  |  |
| --- | --- | --- | --- | --- |
| **Target** | **Fluorochrome** | **Clone** | **Source** | **RRID** |
| CD19 | BUV395 | 1D3 | BD Biosciences | AB_2722495 |
| CD8a | PerCPeFluor710 | 53-6.7 | eBioscience | AB_1834433 |
| CD8a | PerCP/Cy5.5 | 53-6.7 | eBioscience | AB_1107004 |
| SiglecF | PE | E50-2440 | Thermo Fisher Scientific |  |
| Ly6c | eFluor450 | HK1.4 | Thermo Fisher Scientific | AB_10805519 |
| CD11c | FITC | N418 | Thermo Fisher Scientific | AB_464940 |
| GR1 | PE cy7 | RB6-8C5 | Thermo Fisher Scientific | AB_469663 |
| TCRβ | APCeFluor780 | H57-597 | eBioscience | AB_1272173 |
| TCRβ | PE | H57-597 | BD Biosciences | AB_394684 |
| CD4 | Alexa647 | GK1.5 | BioLegend | AB_493372 |
| CD4 | BV421 | GK1.5 | BioLegend | AB_10900241 |
| CD11b | Alexa700 | M1/70 | In house |  |
| Anti-rabbit IgG | Alexa Fluor 647 |  | Life Technologies, A21244 | AB_2535812 |
| Myc |  | D84C12 | Cell Signaling, #5605 |  |

## METHODS

### Generation of Myce mouse

For generating the Myce mouse strain, a 215bp region(chr15:61,418,510-61,418,725) approximately 438,000bp upstream of the *Myc* promoter on chromosome 15 was deleted by CRISPR editing using two single gRNAs with the sequences 5’-ATAGATCATGACCTCCAAAG-3’ and 5’-GACTTTGAACGTTACCTTCT-3’ following the protocol by Aubery et al.^37^. Mice were genotyped using the forward primer 5’-GTTTTATTTCTTCCCCAAGGCAC-3’ and reverse primer 5’-AGCACTGGGTCTCTGAACTG-3’. A PCR product of 394 bp is expected if the deleted allele is present. Deletion was confirmed by gel electrophoresis and next generation sequencing (Supplementary Figure 2B-C). Myce−/− mice were generated on a C57BL/6J background and crossed to C57BL/6J for 4 generations prior to experimental use.

### Mouse immunophenotyping

Spleen, lymph node and thymus of Myce and littermate WT control were harvested into single-cell suspension with red cell lysis and stained with CD19-BUV395 (1D3, BD), TCRβ-PE (H57-597, BD), CD4-BV421 (GK1.5, BioLegend) and CD8a-PerCp/Cy5.5 (53-6.7, eBioscience). Flow cytometry analysis was performed on a BD LSRFortessa X-20 analyser.

### *In vitro* cell culture and activation

B lymphocytes or the A20 cell line were cultured in RPMI 1640 with 2mM GlutaMAX, 50 μM β-mercaptoethanol, and 10% heat-inactivated fetal calf serum (FCS) and stimulated with 25 μg/ml lipopolysaccharide (LPS) (Salmonella typhosa origin, Sigma). CD8 T cells were stimulated using anti-CD3 (10μg/ml plate-bound, clone 145-2C11, WEHI antibody facility), anti-CD28 (2μg/ml, clone 3751, WEHI antibody facility) and human IL-2 (Peprotech). CD8 T cell cultures contained 25μg/ml anti-mouse IL2 (clone S4B6, WEHI antibody facility), which neutralises the activity or mouse but not human IL-2 *in vitro ^38^.* For experiments in which cell division number was tracked, cells were labelled with 10μM CellTrace Violet (CTV, Invitrogen), following the manufacturers protocol.

### Intracellular Myc staining

For intracellular staining of Myc, cells were harvested, centrifuged at 500g at 4°C for 5 mins before resuspension in fixation buffer (0.5% PFA + 0.2% Tween 20 in PBS) and incubation at 4°C for 24hrs. Cells were washed with FACS buffer before pelleting at 500g at 4°C for 5 mins and maintained in FACS buffer at 4°C until staining of all samples within each experiment could be performed simultaneously. For staining, cells were incubated with either anti-Myc (clone D84C12, Cell Signaling) or a rabbit IgG isotype control antibody in PBS-0.1% BSA for 45mins at room temperature. Cells were washed with PBS-0.1% BSA before incubating with anti-rabbit IgG Alexa Fluor 647 (A21244, Life Technologies) in PBS-0.1% BSA, for 45mins at room temperature. Cells were washed and flow cytometry analysis performed.

### Calculation of total cell and cohort number

Total cell numbers were calculated based on the ratio of live cells to counting beads detected by flow cytometry in each sample, after a known number of beads was added prior to analysis. Dead cell exclusion was performed using propidium iodide (0.2μg/ml, Sigma). Cohort number and mean division number over time were calculated based upon CellTrace Violet dilution, using the ‘precursor cohort method’ as previously described^39^.

### Asthma induction and analysis

Allergic lung inflammation was induced in Myce and wildtype mice using intraperitoneal ovalbumin (OVA)/alum sensitization followed by nebulised OVA challenge as previously described^22^. The day after the final challenge, ~200μL of blood was collected into non-heparinised tubes by retro-orbital bleeding, and mice were then sacrificed by CO_2_inhalation after which bronchoalveolar lavage (BAL) was performed (250μL twice with sterile PBS). Flow cytometry was performed fresh on BAL cells using the following staining panel: CD19-BUV395 Clone#1D3, CD8a-PerCPeFluor710 Clone#52-6.7, SiglecF-PE Clone#E50-2440 from BD Pharmigen, Ly6c-eFluor450 Clone#HK1.4, CD11c-FITC Clone #N418, GR1-PECy7 Clone#RB6-8C5, TCRβ-APCeFluor780 Clone #H57-597 from eBioscience, CD4-Alexa647 Clone#GK1.5, CD11b-Alexa700 Clone#M1/70 from WEHI Antibody Facility (Supp Table 3). Flow Cytometry was performed on a BD LSRFortessa X-20 analyser. SYTOX Blue Dead Cell Stain was used to exclude dead cells from analysis and SPHERO Rainbow Beads (BD Biosciences) were used to calculate absolute cell counts.

### Influenza infection and analysis

Myce^+/−^ and wildtype Mice were infected with Influenza A/H3N2/X31 virus at 1×10^4^pfu in 25ul by intranasal insufflation on day 0. On day 8, mice were sacrificed by CO_2_ inhalation after which bronchoalveolar lavage (BAL) was performed (250μL twice with sterile PBS). ~200μL of blood was collected into non-heparinised tubes by retro-orbital bleeding, and mice were then sacrificed by CO_2_inhalation after which bronchoalveolar lavage (BAL) was performed (250μL twice with sterile PBS). Flow cytometry was performed fresh on BAL cells using the following staining panel: CD19-BUV395 Clone#1D3, CD8a-PerCPeFluor710 Clone#52-6.7 from BD Pharmigen, CD11c-FITC Clone #N418, GR1-PECy7 Clone#RB6-8C5, TCRβ-APCeFluor780 Clone #H57-597 from eBioscience, CD4-Alexa647 Clone#GK1.5, CD11b-Alexa700 Clone#M1/70 from WEHI Antibody Facility (Supp Table 3). MHC Class I tetramers PA224-BV421 and NP366-PE (a generous gift from Prof Katherine Kedzierska, The University of Melbourne) were used to detect influenza-specific CD8^+^ T Cells. Flow Cytometry was performed on a BD LSRFortessa X-20 analyser. SYTOX Blue Dead Cell Stain was used to exclude dead cells from analysis and SPHERO Rainbow Beads (BD Biosciences) were used to calculate absolute cell counts.

### *In vitro* transcription of sgRNA

sgRNA used in this study were generated via *in vitro* transcription, as previously described^40^. In brief, transcription template was generated by PCR using Q5 high fidelity DNA polymerase (New England Biolabs) with dNTP (Promega) and an annealing temperature of 58°C. Universal primers as well as a specific primer bearing the sgRNA flanked by T7 promoter sequence and scaffold (as the format T7-sgRNA-scaffold) were used (Supplementary Table 2). PCR products were purified by DNA clean & concentrator-25 (Zymo Research). *In vitro* transcription was performed by incubating 5 μg of transcription template with NTP, pyrophosphatase (ThermoFisher,), RNase Inhibitor (Lucigen) and NxGen T7 RNA polymerase (Lucigen) at 37 °C for 18 hours. Transcription template was then digested by TURBO DNase (Invitrogen) at 37 °C for 45 minutes. The remaining sgRNA was then purified by RNA clean & concentrator-25 (Zymo Research).

### Ribonucleoprotein (RNP) assembly and delivery

For Cas9 RNP, 150 pmol of *in vitro* transcribed sgRNA was incubated with 100 pmol of recombinant Cas9 nuclease (Integrated DNA Technologies) at room temperature for 15 minutes. RNP with 100 pmol of electroporation enhancer (IDT) were subsequently transfected into cells via electroporation. Delivery into A20 cells was performed via 4D-Nucleofector (Lonza) with buffer SF and pulse code FF113.

### Generating and confirming the deletion of putative *Myc* enhancers

Deletion of candidate regions in the A20 cell line genome was performed using electroporation delivered RNP containing paired sgRNAs (Supplementary Table 2). 48 hours post-electroporation single cell clones were sorted into U-bottom 96-well plates and expanded as outlined above. A fraction of the expanded population was lysed in Direct PCR Lysis reagent (Viagen) with proteinase K (Roche). Genotyping was performed directly from the lysates with region targeting primers (Supplementary Table 2) and MyTaq Red Mix (Bioline) followed by gel electrophoresis to reveal biallelic knockout clones. Expected deletions are shown in Supplementary Figure 2.

### Quantitative reverse transcription PCR (qRT-PCR)

RNA was extracted using NucleoSpin RNA Plus (Macherey-Nagel) with gDNA removal, 1 μg of RNA was then reverse transcribed with random hexamer and anchored oligo dT primers using SensiFast cDNA synthesis kit (Bioline). Quantitative PCR was performed on the diluted cDNA using SensiFast probe (Bioline) as per manufacturers’ protocol with β-actin acting as an endogenous reference. Primer and probe sequences are detailed in Supplementary Table 2. Gene expression was normalised to the endogenous control and relative expression was evaluated using ΔΔCt method.

### *In situ* HiC

The *in situ* HiC data is available as GEO series GSE147467. The HiC libraries were processed and differential interactions were assessed as previously described^1^.

### 3C from HiC

Fixation, proximity ligation and DNA precipitation were performed as for *in situ* HiC^41^, except the 0.4mM biotinylated dATP (Life Technologies, 19524-016) in the end blunting step was replaced with 10mM unbiotinylated dATP (NEB, N0440S). The protocol was ceased after DNA precipitation and before DNA shearing. The proximity ligated DNA was then quantified using a Nanodrop (Thermo Fisher Scientific).

The frequency of proximity ligated fragments containing the *Myc* promoter and other downstream regions was then quantified on the BioRad C1000 Thermo cycler using the SensiMix SYBR NOROX kit (Bioline) with 60°C annealing temperature, water or 5ng of DNA per 25ul reaction in duplicate. Primers P1 and P4 (Supplementary Table 2) were used to capture the ligation events between the *Myc* promoter itself. The P3 primer to all other regions numbered sequentially according to their location across the 1Mb upstream of the *Myc* promoter (Supplementary Table 2) were used to quantify interactions between the *Myc* promoter and other regions. Primer #22 targeted the Myce deleted region. Data was plotted as the average raw threshold cycle (Ct) in DNA samples per primer pair minus the average threshold cycle of the paired water controls. Relative to WT was calculated by subtracting the Myce mice Ct value from the associated WT Ct.

### Virtual 4C analysis

The virtual 4C profiles were produced with the diffHic package v1.26.0 in R and then plotted using Graphpad PRISM. The myc promoter was defined with the TxDb.Mmusculus.UCSC.mm10.knownGene package v3.10.0 and by applying the promoters function from the GenomicFeatures package v1.46.2 with upstream□=□2kbp and downstream□=□5kbp. Interactions between the *Myc* promoter and the entire genome were counted across all samples with the connectCounts function from diffHic using regions=myc promoter with a filter set to 0 and the second.region□=□20kbp. The counts per million for each bin were calculated with the cpm function from the edgeR package v3.36.0.

### Visualization of HiC

Normalized contact matrices at a 20 kbp resolution were produced with the HOMER HiC pipeline for visualization. With the summed biological-replicate tag directories, the analyzeHiC function was used with the -balance option. Contact matrices were plotted using the plotHic function from the Sushi R package v1.34.0^42^. The color palette was inferno from the viridisLite package v0.4.0^43^.

### RNA-Sequencing analysis

The RNA-seq data is available as GEO series GSE147496. The RNA-Seq libraries were processed and differential expression was assessed as previously described^1^.

## Acknowledgements

We thank the staff of the core facilities at the Walter and Eliza Hall Institute. This work was supported by grants and fellowships from the Marian and E.H. Flack Fellowship (H.D.C.), the National Health and Medical Research Council of Australia (C.R.K #1125436, T.M.J #1124081, R.S.A #1100451, G.K.S & R.S.A #1158531), and the Australian Research Council (R.S.A. #130100541). This study was made possible through Victorian State Government Operational Infrastructure Support, the Australian Government NHMRC Independent Research Institute Infrastructure Support scheme, and the Australian Cancer Research Fund. The generation of Myce −/− mice used in this study was supported by Phenomics Australia and the Australian Government through the National Collaborative Research Infrastructure Strategy program.

## Author contributions

W.F.C., M.R., N.I., C.A., C.R.K., J.R.G., A.K.W. and T.M.J. designed and conducted experiments. A.J.K. generated the Myce mice. H.D.C. and G.K.S. performed computational analyses. T.M.J., and R.S.A. conceived the study and wrote the manuscript with assistance from H.D.C.

## Competing interests

The authors declare no competing interests

## Code availability

DiffHic and edgeR, which are freely available from the Bioconductor repository (https://bioconductor.org/packages/release/bioc/html/diffHic.html and https://bioconductor.org/packages/release/bioc/html/edgeR.html, respectively) were used in this study. Versions used for this manuscript were: diffHic v1.26.019, edgeR package v3.36.0^44^, Sushi R package v1.22.0^45^, HOMER v4.11^46^.

## Notes

### Competing Interest Statement

The authors have declared no competing interest.

### Summary of Updates

New figures have been added because they were not ok in the first submission

## References

1 Chan, W. F. et al. Pre-mitotic genome re-organisation bookends the B cell differentiation process. Nat Commun 12, 1344 (2021). https://doi.org:10.1038/s41467-021-21536-2

2 Zan, H. & Casali, P. Epigenetics of Peripheral B-Cell Differentiation and the Antibody Response. Front Immunol 6, 631 (2015). https://doi.org:10.3389/fimmu.2015.00631

3 Schmidl, C., Delacher, M., Huehn, J. & Feuerer, M. Epigenetic mechanisms regulating T-cell responses. J Allergy Clin Immunol 142, 728–743 (2018). https://doi.org:10.1016/j.jaci.2018.07.014

4 Marchingo, J. M., Sinclair, L. V., Howden, A. J. & Cantrell, D. A. Quantitative analysis of how Myc controls T cell proteomes and metabolic pathways during T cell activation. Elife 9. https://doi.org:10.7554/eLife.53725

5 Assmann, N. & Finlay, D. K. Metabolic regulation of immune responses: therapeutic opportunities. J Clin Invest 126, 2031–2039 (2016). https://doi.org:10.1172/JCI83005

6 Liu, J. & Levens, D. Making myc. Curr Top Microbiol Immunol 302, 1–32 (2006). https://doi.org:10.1007/3-540-32952-8_1

7 de Alboran, I. M. et al. Analysis of C-MYC function in normal cells via conditional gene-targeted mutation. Immunity 14, 45–55 (2001). https://doi.org:10.1016/s1074-7613(01)00088-7

8 Wang, R. et al. The transcription factor Myc controls metabolic reprogramming upon T lymphocyte activation. Immunity 35, 871–882 (2011). https://doi.org:10.1016/j.immuni.2011.09.021

9 Conacci-Sorrell, M., McFerrin, L. & Eisenman, R. N. An overview of MYC and its interactome. Cold Spring Harb Perspect Med 4, a014357 (2014). https://doi.org:10.1101/cshperspect.a014357

10 Bouchard, C., Staller, P. & Eilers, M. Control of cell proliferation by Myc. Trends Cell Biol 8, 202–206 (1998). https://doi.org:10.1016/s0962-8924(98)01251-3

11 Lancho, O. & Herranz, D. The MYC Enhancer-ome: Long-Range Transcriptional Regulation of MYC in Cancer. Trends Cancer 4, 810–822 (2018). https://doi.org:10.1016/j.trecan.2018.10.003

12 Vervoorts, J., Lüscher-Firzlaff, J. & Lüscher, B. The ins and outs of MYC regulation by posttranslational mechanisms. J Biol Chem 281, 34725–34729 (2006). https://doi.org:10.1074/jbc.R600017200

13 Banerji, J., Rusconi, S. & Schaffner, W. Expression of a beta-globin gene is enhanced by remote SV40 DNA sequences. Cell 27, 299–308 (1981). https://doi.org:10.1016/0092-8674(81)90413-x

14 Corcoran, L. M., Adams, J. M., Dunn, A. R. & Cory, S. Murine T lymphomas in which the cellular myc oncogene has been activated by retroviral insertion. Cell 37, 113–122 (1984). https://doi.org:10.1016/0092-8674(84)90306-4

15 Cory, S. et al. Activation of the c-myc oncogene in B and T lymphoid tumors. Curr Top Microbiol Immunol 113, 161–165 (1984). https://doi.org:10.1007/978-3-642-69860-6_27

16 Dekker, J., Rippe, K., Dekker, M. & Kleckner, N. Capturing chromosome conformation. Science 295, 1306–1311 (2002). https://doi.org:10.1126/science.1067799

17 Sati, S. & Cavalli, G. Chromosome conformation capture technologies and their impact in understanding genome function. Chromosoma 126, 33–44 (2017). https://doi.org:10.1007/s00412-016-0593-6

18 Cho, S. W. et al. Promoter of lncRNA Gene PVT1 Is a Tumor-Suppressor DNA Boundary Element. Cell 173, 1398–1412.e1322 (2018). https://doi.org:10.1016/j.cell.2018.03.068

19 Lun, A. T. & Smyth, G. K. diffHic: a Bioconductor package to detect differential genomic interactions in Hi-C data. BMC Bioinformatics 16, 258 (2015). https://doi.org:10.1186/s12859-015-0683-0

20 Yoshida, H. et al. The cis-Regulatory Atlas of the Mouse Immune System. Cell 176, 897–912.e820 (2019). https://doi.org:10.1016/j.cell.2018.12.036

21 Mahajan, V. S. et al. B1a and B2 cells are characterized by distinct CpG modification states at DNMT3A-maintained enhancers. Nat Commun 12, 2208 (2021). https://doi.org:10.1038/s41467-021-22458-9

22 Keenan, C. R. et al. Polycomb repressive complex 2 is a critical mediator of allergic inflammation. JCI Insight 4. https://doi.org:10.1172/jci.insight.127745

23 Kvon, E. Z., Waymack, R., Gad, M. & Wunderlich, Z. Enhancer redundancy in development and disease. Nat Rev Genet 22, 324–336 (2021). https://doi.org:10.1038/s41576-020-00311-x

24 Twell, D., Yamaguchi, J., Wing, R. A., Ushiba, J. & McCormick, S. Promoter analysis of genes that are coordinately expressed during pollen development reveals pollen-specific enhancer sequences and shared regulatory elements. Genes Dev 5, 496–507 (1991). https://doi.org:10.1101/gad.5.3.496

25 Camprodón, F. J. & Castelli-Gair, J. E. Ultrabithorax protein expression in breakpoint mutants: localization of single, co-operative and redundant cis regulatory elements. Rouxs Arch Dev Biol 203, 411–421 (1994). https://doi.org:10.1007/BF00188690

26 Ertzer, R. et al. Cooperation of sonic hedgehog enhancers in midline expression. Dev Biol 301, 578–589 (2007). https://doi.org:10.1016/j.ydbio.2006.11.004

27 Grosveld, F., van Assendelft, G. B., Greaves, D. R. & Kollias, G. Position-independent, high-level expression of the human beta-globin gene in transgenic mice. Cell 51, 975–985 (1987). https://doi.org:10.1016/0092-8674(87)90584-8

28 Osterwalder, M. et al. Enhancer redundancy provides phenotypic robustness in mammalian development. Nature 554, 239–243 (2018). https://doi.org:10.1038/nature25461

29 Wang, X. & Goldstein, D. B. Enhancer Domains Predict Gene Pathogenicity and Inform Gene Discovery in Complex Disease. Am J Hum Genet 106, 215–233 (2020). https://doi.org:10.1016/j.ajhg.2020.01.012

30 Bahr, C. et al. A Myc enhancer cluster regulates normal and leukaemic haematopoietic stem cell hierarchies. Nature 553, 515–520 (2018). https://doi.org:10.1038/nature25193

31 Fulco, C. P. et al. Systematic mapping of functional enhancer-promoter connections with CRISPR interference. Science 354, 769–773 (2016). https://doi.org:10.1126/science.aag2445

32 Bothma, J. P. et al. Enhancer additivity and non-additivity are determined by enhancer strength in the Drosophila embryo. Elife 4. https://doi.org:10.7554/eLife.07956

33 Lam, D. D. et al. Partially redundant enhancers cooperatively maintain Mammalian pomc expression above a critical functional threshold. PLoS Genet 11, e1004935 (2015). https://doi.org:10.1371/journal.pgen.1004935

34 La Rosa, F. A., Pierce, J. W. & Sonenshein, G. E. Differential regulation of the c-myc oncogene promoter by the NF-kappa B rel family of transcription factors. Mol Cell Biol 14, 1039–1044 (1994). https://doi.org:10.1128/mcb.14.2.1039-1044.1994

35 Chen, H. et al. Dynamic interplay between enhancer-promoter topology and gene activity. Nat Genet 50, 1296–1303 (2018). https://doi.org:10.1038/s41588-018-0175-z

36 Johanson, T. M., Keenan, C. R. & Allan, R. S. Shedding Structured Light on Molecular Immunity: The Past, Present and Future of Immune Cell Super Resolution Microscopy. Front Immunol 12, 754200 (2021). https://doi.org:10.3389/fimmu.2021.754200

37 Aubrey, B. J. et al. An inducible lentiviral guide RNA platform enables the identification of tumor-essential genes and tumor-promoting mutations in vivo. Cell Rep 10, 1422–1432 (2015). https://doi.org:10.1016/j.celrep.2015.02.002

38 Deenick, E. K., Gett, A. V. & Hodgkin, P. D. Stochastic model of T cell proliferation: a calculus revealing IL-2 regulation of precursor frequencies, cell cycle time, and survival. J Immunol 170, 4963–4972 (2003). https://doi.org:10.4049/jimmunol.170.10.4963

39 Marchingo, J. M. et al. T cell signaling. Antigen affinity, costimulation, and cytokine inputs sum linearly to amplify T cell expansion. Science 346, 1123–1127 (2014). https://doi.org:10.1126/science.1260044

40 Chan, W. F. et al. Identification and characterization of the long noncoding RNA Dreg1 as a novel regulator of Gata3. Immunol Cell Biol 99, 323–332 (2021). https://doi.org:10.1111/imcb.12408

41 Rao, S. S. et al. A 3D map of the human genome at kilobase resolution reveals principles of chromatin looping. Cell 159, 1665–1680 (2014). https://doi.org:10.1016/j.cell.2014.11.021

42 Phanstiel, D. H., Boyle, A. P., Araya, C. L. & Snyder, M. P. Sushi.R: flexible, quantitative and integrative genomic visualizations for publication-quality multi-panel figures. Bioinformatics 30, 2808–2810 (2014). https://doi.org:10.1093/bioinformatics/btu379

43 Garnier, S. et al. (2022).

44 Robinson, M. D., McCarthy, D. J. & Smyth, G. K. edgeR: a Bioconductor package for differential expression analysis of digital gene expression data. Bioinformatics 26, 139–140 (2010). https://doi.org:10.1093/bioinformatics/btp616

45 Sushi: Tools for visualizing genomics data (2015).

46 Heinz, S. et al. Simple combinations of lineage-determining transcription factors prime cis-regulatory elements required for macrophage and B cell identities. Mol Cell 38, 576–589 (2010). https://doi.org:10.1016/j.molcel.2010.05.004

